# From introduction to eradication: reconstructing population size and removal history of an invasive species

**DOI:** 10.64898/2026.07.29.737110

**Authors:** Keita Fukasawa, Takuma Sato, Takamichi Jogahara, Tomonori Kawamoto, Takahiro Morosawa, Takuma Hashimoto, Masaki Asano, Tamotsu Matsuda, Yoshihito Goto, Shin Hosokawa, Katsushi Nakata, Ryoji Fukuhara, Nobuo Ishii, Yuya Watari, Ken Ishida, Fumio Yamada, Shintaro Abe

**Affiliations:** Biodiversity Division, National Institute for Environmental Studies, 16-2 Onogawa, Tsukuba, Ibaraki 305-8506, Japan; Department of Law, Economics and Management, Okinawa University, 555 Kokuba, Naha, Okinawa 902-8521, Japan; Regional Environment Office in Hokkaido, Ministry of the Environment, Kita 8 Nishi 2, Kita-ku, Sapporo, Hokkaido 060-0808, Japan; Japan Wildlife Research Center, 3-3-7 Kotobashi, Sumida-ku, Tokyo 130-8606, Japan; Wildlife Management Center, Tokyo University of Agriculture and Technology, 3-5-8 Saiwaicho, Fuchu, Tokyo 183-8509, Japan; Nansei Environmental Laboratory Co., Ltd., 4-4 Agarizaki, Nishihara, Okinawa 903-0105, Japan; Tokyo Woman’s Christian University, 2-6-1 Zempukuji, Suginami-ku, Tokyo 167-8585, Japan; Department of Wildlife Biology, Forestry and Forest Products Research Institute, 1 Matsunosato, Tsukuba, Ibaraki, 305-8687, Japan; No affiliation; Amami Rabbit Museum QuruGuru, Mahoroba Park, 502 Ongachi, Yamato, Kagoshima 894-3104, Japan; Amami Wildlife Conservation Center, Ministry of the Environment, 551 Ongachi, Yamato, Kagoshima 894-3104, Japan

**Author notes:** Corresponding author: Keita Fukasawa.

**Keywords:** abundance, catchability, eradication, hierarchical Bayesian model, invasive alien species, population size, state-space model

## Abstract

1. Understanding the processes underlying successful eradication of invasive species is essential for achieving global island conservation goals. Despite the widespread availability of capture records from eradication programs, modeling frameworks that utilize these datasets to elucidate spatio-temporal population dynamics remain underdeveloped.
2. In this study, we reconstructed the spatio-temporal population dynamics of the small Indian mongoose on Amami-Oshima Island (712 km²), Japan, where the species was introduced in 1979 and officially declared eradicated in 2024 after more than 30 years of systematic removal. We integrated introduction records, capture data, and monitoring data using a hierarchical harvest-based model (HBM). To evaluate the model’s capacity to support management decisions and assess eradication success, we conducted retrospective analyses and compared estimated eradication probabilities with those obtained from a rapid eradication assessment (REA; Samaniego-Herrera et al., 2013).
3. The estimated population size (before reproduction) peaked at 5,449 individuals (95% CI: 4,703, 6,175) in 2000 and subsequently declined almost monotonically. The maximum invaded area was 547.78 km² (posterior median, 95% CI: 496.47, 566.04) in 2009, indicating that the removal program successfully prevented island-wide expansion. Retrospective analyses showed that population estimates remained within the 95% credible intervals of the full dataset estimates, demonstrating temporal consistency. Eradication probabilities estimated by the HBM were substantially higher than those from the REA, highlighting the sensitivity of estimates to fine-scale heterogeneity in detection processes.
4. Synthesis and applications: Hierarchical HBMs provide a powerful framework for reconstructing, predicting, and evaluating invasive species eradication dynamics. Being aware of the limitations for application to eradication evaluations, HBMs can support adaptive management in long-term eradication programs and improve our understanding of the mechanisms underlying successful eradication.

## Introduction

Invasive alien species are a major threat to global biodiversity and human well-being (Roy et al., 2024). Biodiversity loss on islands is disproportionately rapid compared to that on the mainland, with invasive species identified as a primary driver of extinction (Spatz et al., 2017). Eradication of invasive species is therefore a promising strategy for restoring island ecosystems and biodiversity (Jones et al., 2016). Although many island eradication campaigns have been successful (DIISE, 2019), eradication on large islands remains particularly challenging (Capizzi, 2020).

Eradication efforts on large islands typically require long-term, often multi-decadal, commitments (Fukasawa, Hashimoto, et al., 2013; Legge et al., 2018). Sustained investment and adaptive management are essential to increase the likelihood of success. To support effective adaptive decision-making, it is crucial to evaluate current population status, control efficiency, and long-term outcomes of management actions (Nishimoto et al., 2020). However, estimating these quantities is notoriously difficult because direct observations of population size are rarely available for invasive species, which are often cryptic.

Fukasawa et al. (2013) developed a hierarchical harvest-based model (HBM) that integrates information on introduced population size, capture data, and underlying population dynamics based on a surplus-production framework commonly used in fisheries stock assessment (Hilborn & Walters, 1992). While this approach enables the estimation of capture efficiency and eradication feasibility conditional on effort, it does not explicitly account for spatial population structure. This limitation is critical when assessing eradication probability, because such estimates are sensitive to assumptions about spatial processes (Samaniego-Herrera et al., 2013).

Diffusive spread is a fundamental characteristic of many organisms and is explicitly incorporated in the rapid eradication assessment (REA; Samaniego-Herrera et al., 2013), a widely used spatially explicit simulation framework for evaluating eradication probability. Few studies have incorporated diffusion processes into HBMs (Osada et al., 2019), and their potential for predicting management outcomes of invasive species remains insufficiently explored. In particular, the stability of parameter estimates, predictive performance (Carvalho et al., 2021), and behavior of these models in the context of eradication probability assessment require further evaluation.

The eradication of the small Indian mongoose (*Urva auropunctata*, Fig. 1a) on Amami-Oshima Island (712km^2^), Japan, provides a rare example of a large-scale, long-term eradication project. The species was introduced in 1979 and caused severe declines in endemic fauna (Yamada & Sugimura, 2004; Watari et al., 2008). Intensive and systematic removal using traps began in 1993, leading to a population decline of mongoose (Fukasawa, Hashimoto, et al., 2013) and recovery of native species (Fukasawa, Miyashita, et al., 2013; Watari et al., 2013). Sniffer dogs for searching individuals and scats had been applied since 2011 and 2014, respectively, and proved effective in detecting remaining mongooses (Fukuhara et al., 2010; Mitani et al., 2014). No individuals have been detected since the last capture in 2018, and eradication was officially declared in 2024 (https://www.env.go.jp/en/press/press_03205.html). The capture dataset is exceptionally comprehensive, including annual capture numbers, trap effort, and spatial information since 2000, enabling the development of spatially explicit models spanning the entire invasion-to-eradication process.

**Fig. 1.**
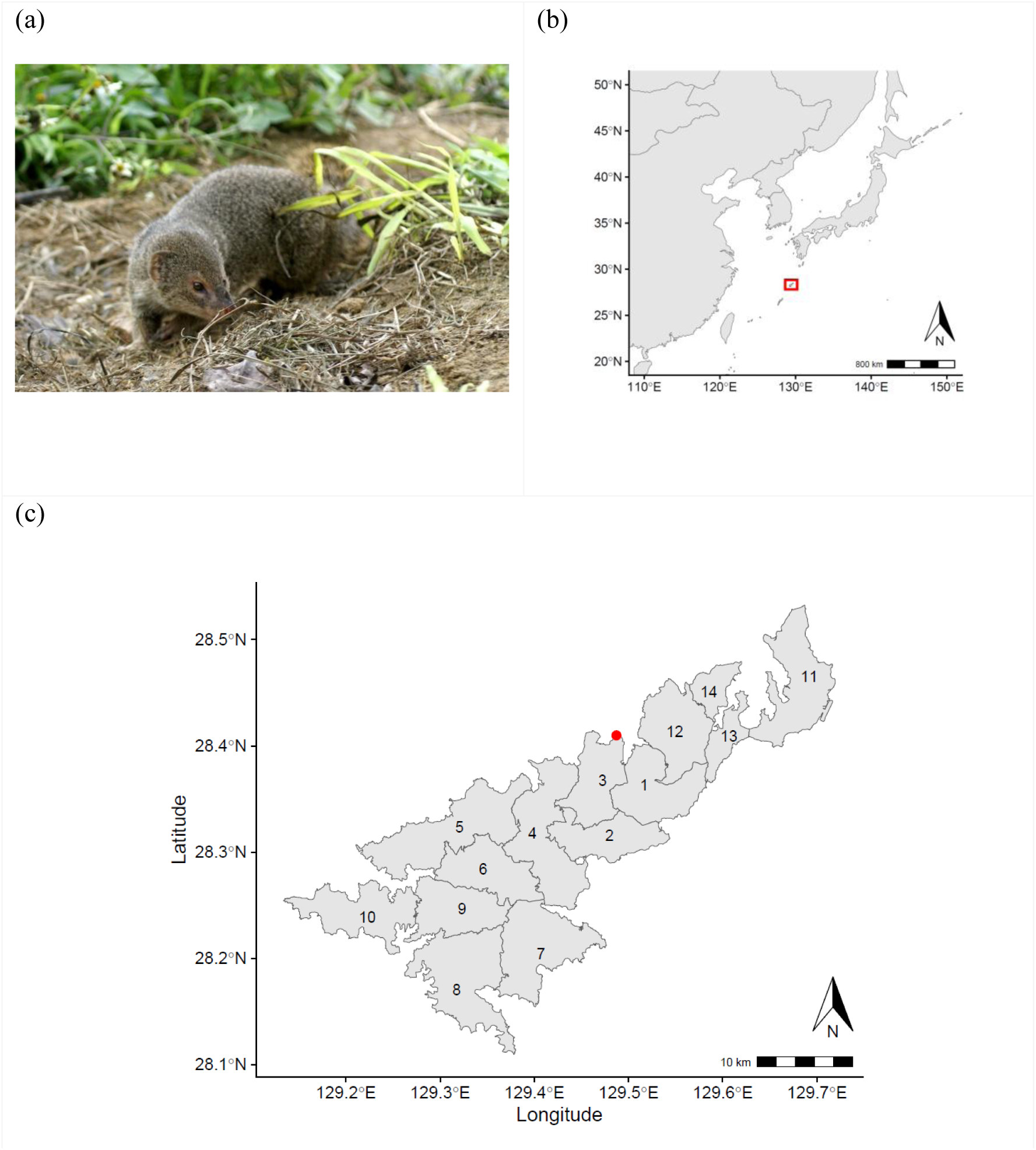
(a) small Indian mongoose in Amami-Oshima Island, (b) location of Amami-Oshima island, (c) the mongoose release point (red dot) and the 14 mongoose management units of Amami-Oshima Island. The id number of units were shown on the map.

In this study, we aimed to reconstruct the 46-year spatio-temporal dynamics of mongoose population size and capture processes on Amami-Oshima Island using a hierarchical HBM that incorporates diffusion among spatial units. To evaluate model performance, we conducted retrospective analyses and hindcast cross-validation (Carvalho et al., 2021). These analyses allowed us to assess the stability of population estimates with the addition of new data and the predictive accuracy of the model for unobserved capture and monitoring data. Furthermore, we estimated eradication probability following the final detection in 2018 using the hierarchical HBM and compared the results with those obtained from the REA framework to evaluate applicability of the population size reconstruction to eradication confirmation.

## Materials and Methods

### Study site

Amami-Oshima Island (712 km²) is located in the Ryukyu Archipelago of southwestern Japan (Fig. 1b) and is characterized by a humid subtropical climate, with a mean annual temperature of 21.8 °C and annual precipitation of 2,935.7 mm (for 1991∼2020, Japan Meteorological Agency). The island is dominated by evergreen broadleaf forests interspersed with secondary habitats that support numerous endemic and endangered species.

The small Indian mongoose was introduced to Amami-Oshima Island in 1979 as a biological control agent targeting a native venomous snake (*Protobothrops flavoviridis*). Thirty individuals were released in a forested area near Amami City (Yamada & Sugimura, 2004) (Fig. 1c). As the population expanded across the island, its ecological impacts on native fauna rapidly became evident. The species posed a major threat to ground-dwelling native birds, mammals, amphibians, and reptiles that had evolved in the absence of mammalian predators (Yamada & Sugimura, 2004; Watari et al., 2008).

Initial control efforts in the 1980s relied on farmer-led trapping, which evolved into a bounty program coordinated by the local government in 1993. In 2000, a formal eradication program led by Ministry of the Environment was launched. The transition from the bounty system to a fully government-operated removal program was completed in 2005, when captures came to be conducted exclusively by trained personnel (the “Amami Mongoose Busters”).

Live traps were used throughout the program, and successive generations of kill traps were introduced from 2003 onward to improve efficiency and reduce bycatch. Dog-assisted capture methods were introduced in 2011, in which trained sniffer dogs (e.g., Terrier and Pointer breeds) were used to locate individual mongooses, allowing handlers to directly capture detected individuals (Mitani et al., 2014). In addition, scat-detection dogs (e.g., German Shepherds) were deployed from 2014 to monitor remaining individuals. These surveys provided a relative abundance index based on the number of scats detected per unit distance.

To monitor the progress of eradication with high spatial resolution and to allocate the trapping effort in space efficiently, the island was divided into 14 management units in 2013 (Fig. 1c). In the HBM, all the observations were aggregated into the management units. The number of captured individuals was recorded annually. Capture effort and trap locations were partially recorded from 1996 and fully recorded from 2001 onward. Trap effort was standardized as corrected trap-days (CTD) to account for differences among trap types and checking intervals (Fukasawa, Hashimoto, et al., 2013). CTD increased substantially after 2001, while catch per unit effort declined (Fig. S1a). A limited application of diphacinone bait was conducted in 2017 and 2018 over approximately 0.1 km^2^; however, this was not included in our model due to its small spatial and temporal extent. All population indices declined over time, and no captures or detections have been recorded since the last capture in 2018 (Fig. S1abc).

### Hierarchical harvest-based models with spatial structure

To reconstruct the spatio-temporal dynamics of population size, we integrated introduction records, capture data from traps and dog-assisted methods, and scat monitoring data. Data with spatial information were aggregated by Japanese fiscal year (April–March; hereafter “year”) and across the 14 management units. The introduction record defined the initial population distribution in 1979: 30 individuals were assigned to the release unit (unit 3; Fig. 1c) and zero to all other units. Capture data consisted of the number of captured individuals and corresponding effort (CTD for traps and search distance [km] for dogs). For earlier records (1979–1999), which lacked spatial and effort information, only total annual captures were available.

We formulated the model within a hierarchical state-space framework consisting of a process model, observation model, and prior distributions. The process model describes the temporal dynamics of population size across spatial units, accounting for intrinsic population growth, dispersal, and removal through capture.

The spatial surplus-production component captures how population size changes due to local reproduction, movement among neighboring units, and removal by capture. In particular, dispersal was modeled as a diffusion process, in which individuals tend to move from high-density areas to low-density areas. Following Fick’s first law of diffusion (Okubo & Levin, 2001), the flux between adjacent units is proportional to the density gradient, and the flow rate is proportional to the length of the shared boundary between units. Let **N***_t_* = (*N_it_*) denote the population size in each unit before population growth in year *t*, and **S***_t_* = (S*_it_*) denote the population size after growth. The process model is defined as:

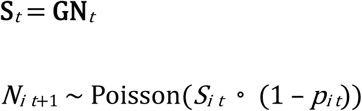

where *p_it_* is the total capture probability in unit *i* and year *t*. The matrix **G** represents population growth and dispersal and is defined as:

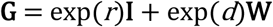

where *r* is the logarithm of (1 + the intrinsic population growth rate), and *d* is the logarithm of the dispersal coefficient. The connectivity matrix **W** = (*w_ij_*) describes dispersal among neighboring units based on shared boundary length. For *i* ≠ *j*, *w_ij_* is *b_ij_* / *A_i_* if units *i* and *j* are adjacent (where *b_ij_* (*= b_ji_*) is the shared boundary length and *A_i_* is the area of unit *i*), and *w_ij_* is zero otherwise. Diagonal elements are defined as *w_ii_* = −Σ_{*j* ≠ *i*}_ *w_ij_* so that each row sums to zero.

Capture processes were modeled using a multi-gear extension of the Weibull catchability framework, allowing for heterogeneous catchability among individuals and across space and time (Fukasawa, Hashimoto, et al., 2013). The total capture probability is given by:

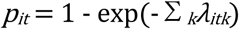

where *λ_itk_* is the capture intensity associated with gear *k*. We considered three types of capture processes: (1) traps with effort data, (2) dog-assisted capture, and (3) trap data without effort information:

Trap (*k* = 1):

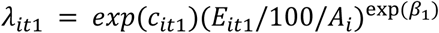

Dog-assisted capture (*k* = 2):

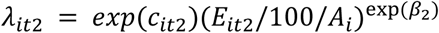

Trap without effort information (*k* = 3):

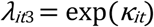

The parameters *c_itk_* represent catchability and were modeled as spatio-temporal random effects:

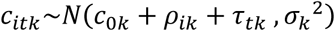

where *c*_0*k*_ is the global mean, and *ρ_ik_* and *τ_tk_* represent spatial and temporal random effects following zero mean normal distribution, respectively. The parameters *β_k_* control the nonlinearity of the relationship between capture effort and capture intensity driven by individual-level heterogeneity in catchability (Fukasawa, Hashimoto, et al., 2013). For trap data without effort information, we modeled *κ_it_* as:

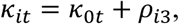

where *κ*_0*t*_ is a year-specific parameter accounting for variation in unknown effort, and *ρ_i_*_3_ represents spatial random effects following zero mean normal distribution.

The capture probability attributable to each gear was defined by partitioning the total capture probability:

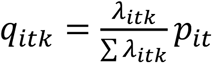

The number of captured individuals for each gear was assumed to follow a Poisson distribution:

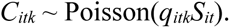

For trap data lacking spatial and effort information, the likelihood was defined at the aggregated annual level:

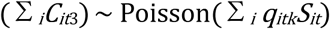

which is consistent with the unit-level formulation due to reproducible property of Poisson distribution.

Scat detections by dogs were treated as an index of population density. The number of detected scats, *Y_it_*, was modelled as:

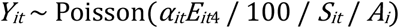

where *E_it_*_4_ denotes the survey effort of scat-detection dogs (measured as search distance in km), and, *α_it_* represents the detection efficiency. To account for spatio-temporal variation in detection efficiency, *α_it_* was modeled as a random effect:

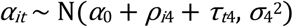

which follows the same structure as the catchability coefficients *c_itk_*, with spatial and temporal random effects capturing heterogeneity in detection processes.

Parameter estimation was conducted within a Bayesian framework using JAGS 4.0.3. Unbounded parameters were assigned Normal priors with mean 0 and variance 25, while variance parameters were assigned standard half-Normal priors. Posterior samples were obtained using 10 parallel Markov chain Monte Carlo (MCMC) chains, each run for 2,000,000 iterations including 1,000,000 burn-in, with thinning every 1,000 iterations, yielding 10,000 posterior samples in total. Convergence was assessed using rank-normalized *R̂* statistics (Vehtari et al., 2021) and visual inspection of the chains.

### Evaluation of Model Consistency

We evaluated model consistency in terms of robustness to data addition and predictive accuracy. To assess robustness, we conducted a retrospective analysis in which observations from the terminal year were sequentially removed backward (“peel”), and population trajectories were re-estimated for each truncated dataset. Standard metrics such as mean absolute scaled error (MASE) are commonly used in fisheries stock assessments (Carvalho et al., 2021), but are not suitable in this context because they become unstable when population size approaches zero, as expected near eradication. Instead, we evaluated consistency by examining whether retrospective point estimates (posterior medians) fell within the 95% credible intervals of the estimates obtained using the full dataset.

Predictive performance was assessed using hindcast cross-validation. For each peel, we generated one-step-ahead predictions for observed data, including the number of captures by traps, the number of dog-assisted captures, and the number of detected scats. Model performance was evaluated by examining whether observed values fell within the corresponding 95% predictive intervals.

### Comparison of eradication probability with rapid eradication assessment

We estimated annual eradication probabilities following the last detection in 2018 using the hierarchical HBM and compared them with estimates obtained from REA (Samaniego-Herrera et al., 2013; Russell et al., 2017). In the hierarchical HBM, the eradication probability in year t is defined as:

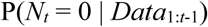

i.e., the probability that the population size equals zero in year *t* conditional on all observations up to year *t* − 1. This quantity was directly computed from posterior samples.

In the REA framework, eradication probability conditional on no detection is derived using Bayes’ theorem:

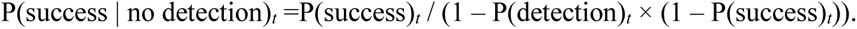

where P(success)*ₜ*denotes the prior probability that eradication has been achieved by year *t*, and P(detection)*ₜ* is the probability of detecting the species in year *t* if it is still present. Detection probability was estimated using spatially explicit individual-based simulations that incorporate reproduction, dispersal, and detection processes, assuming the persistence of a single reproductive individual after the final capture. Model parameters for the simulations were obtained from multiple sources: population growth rate from the hierarchical HBM, dispersal distance derived from historical invasion speed, and detectability and home range size from a spatially explicit capture–recapture (SECR) model newly estimated in this study (Borchers & Efford, 2008). Details in the simulation model were shown in the Appendix S2.

The prior eradication probability in 2019 was initialized using the estimate from the hierarchical HBM, which is the first year in which eradication probability can be defined after the final detection. In subsequent years, eradication probability was updated iteratively based on the absence of detections. The prior probability of eradication in year *t* was calculated as (Samaniego-Herrera et al., 2013):

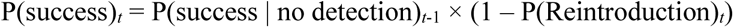

Because intentional translocation of mongooses is strictly prohibited by law in Japan and no cases of accidental introduction across the sea have been reported, the probability of reintroduction was assumed to be negligible and set to zero.

## Results

Using the hierarchical HBM, we reconstructed the full trajectory of population size from introduction to eradication. The posterior median population size peaked at 5,449 individuals (95% CI: 4,703, 6,175) in 2000 and subsequently declined to 125 individuals (95% CI: 104, 149) in 2013 (Fig. 2a). Spatial patterns of population density consistently showed higher densities near the release point across time (Fig. 2b–d; Fig. S2). The total invaded area, Σ*_i_* I(*N_it_* > 0)*A_i_*, reached a maximum of 547.78km^2^ (posterior median, 95%CI: 496.47, 566.04) in 2009. This indicates that the removal program prevented the species from spreading across the entire island, although the spatial range continued to expand even after population size had peaked (Fig. S2, Fig. S3).

**Fig. 2.**
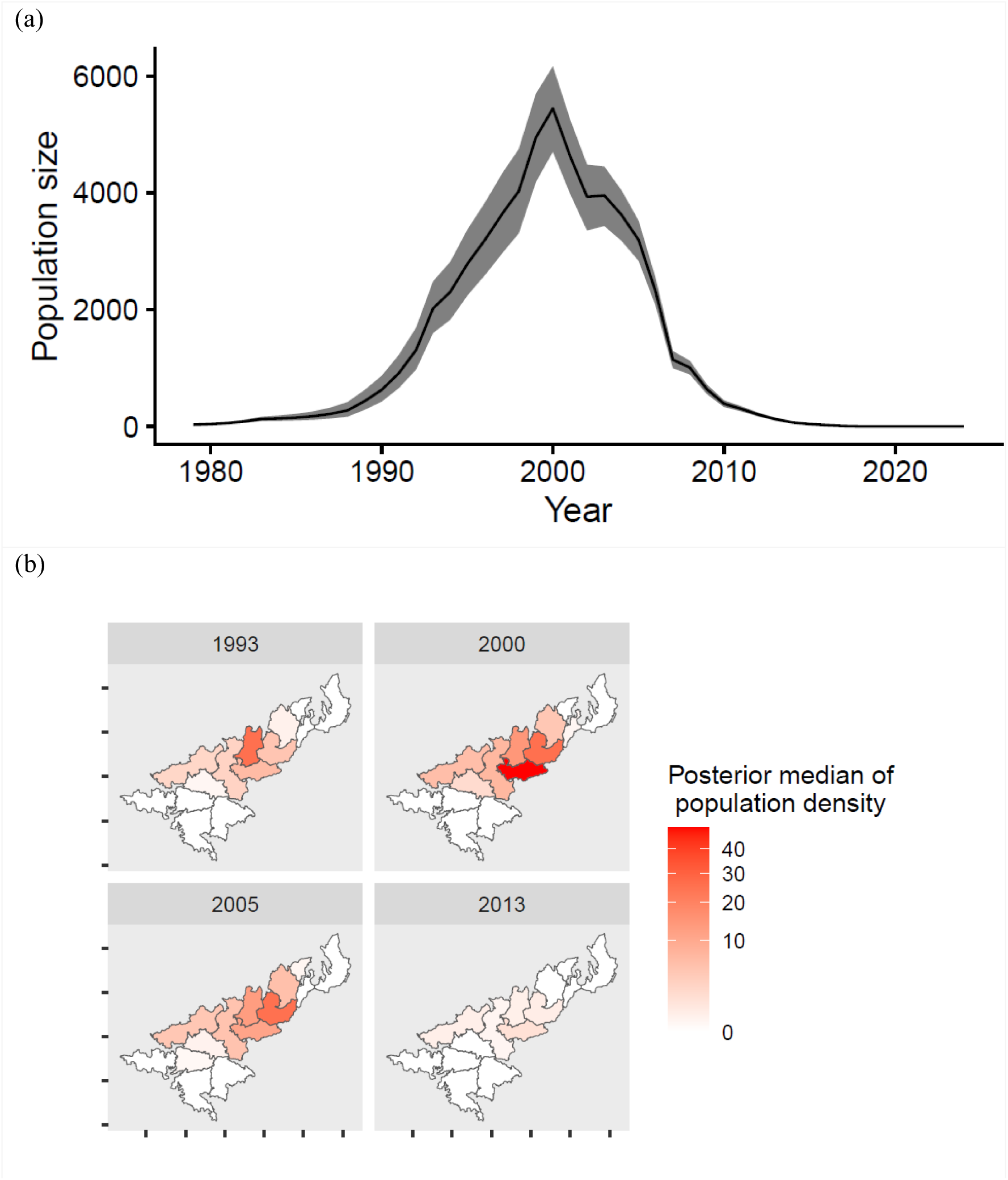
(a) Posterior median and 95% CI of population size, and (b) maps of population density (/km^2^). See Fig. S2 for maps of all the years.

Capture probability increased substantially over time until the last capture in 2018 (Fig. 3a). Although the immediacies of the responses varied, marked increases were observed following key management interventions, including the introduction of the government-led bounty program in 1993, initiation of the eradication program in 2000, the introduction of kill traps in 2003, and the adoption of dog-assisted capture methods in 2011. The spatial extent of capture pressure also expanded over time (Fig. 3b; Fig. S4).

**Fig. 3.**
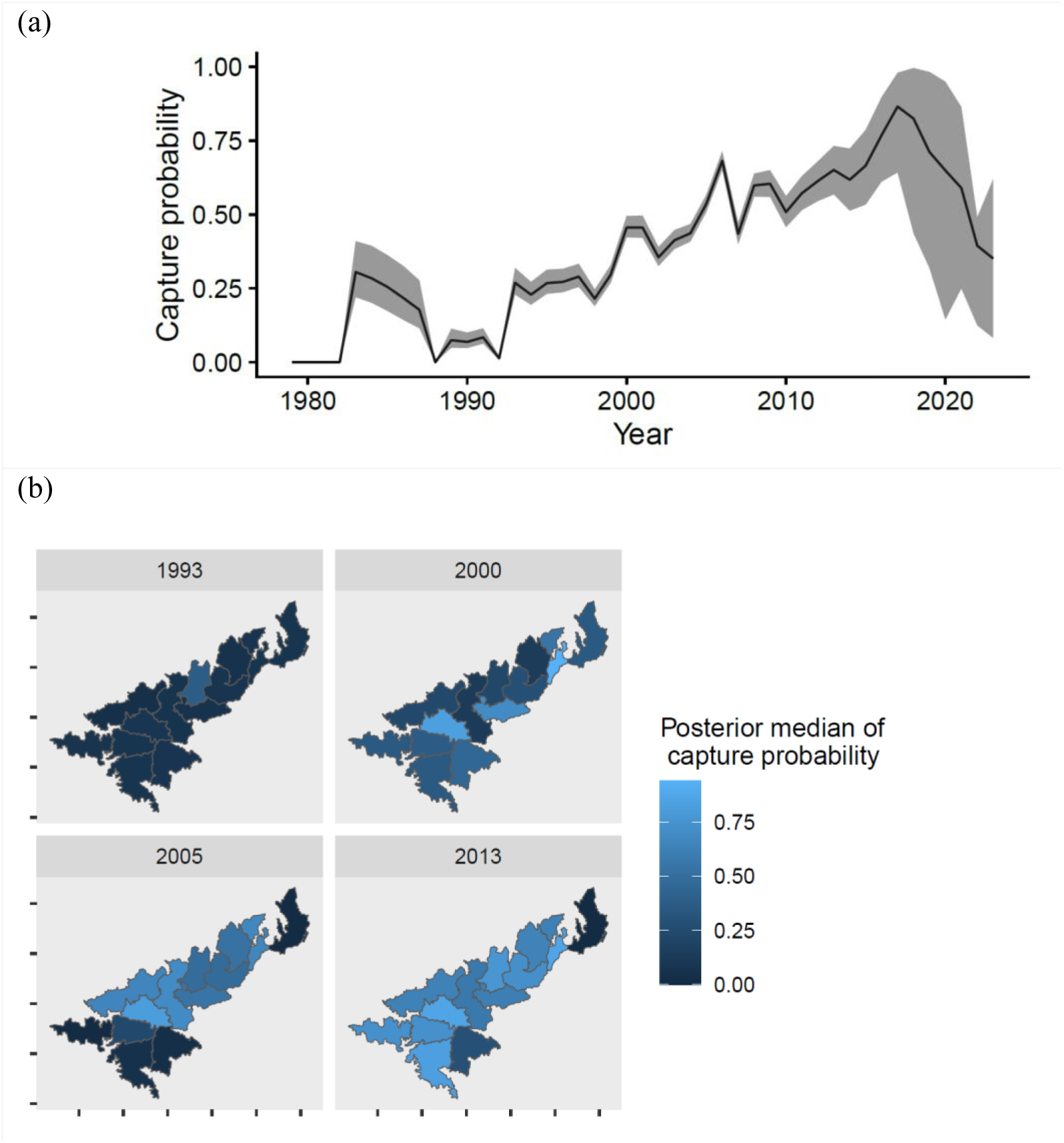
(a) Estimated total capture probability, *p_it_*, indicating capture pressure linked to capture efforts. The solid line and shadow indicate the posterior median and 95% CI, respectively. (b) maps of capture probability. See Fig. S4 for maps of all the years..

Retrospective analysis indicated that population size estimates were stable with respect to the addition of new data. For all peels examined, both total and unit-level population estimates for all years fell within the 95% credible intervals of the estimates based on the full dataset (Fig. 4a; Fig. S5). Results of the hindcast cross-validation were largely consistent with observed data, except for some deviations shortly after the introduction of dog-assisted surveys (Fig. 4b–d; Fig. S6). Specifically, the number of dog-assisted captures in two spatial units in 2015, as well as the number of detected scats in unit 5 in 2016, fell outside the 95% predictive intervals of the one-step-ahead predictions.

**Fig. 4.**
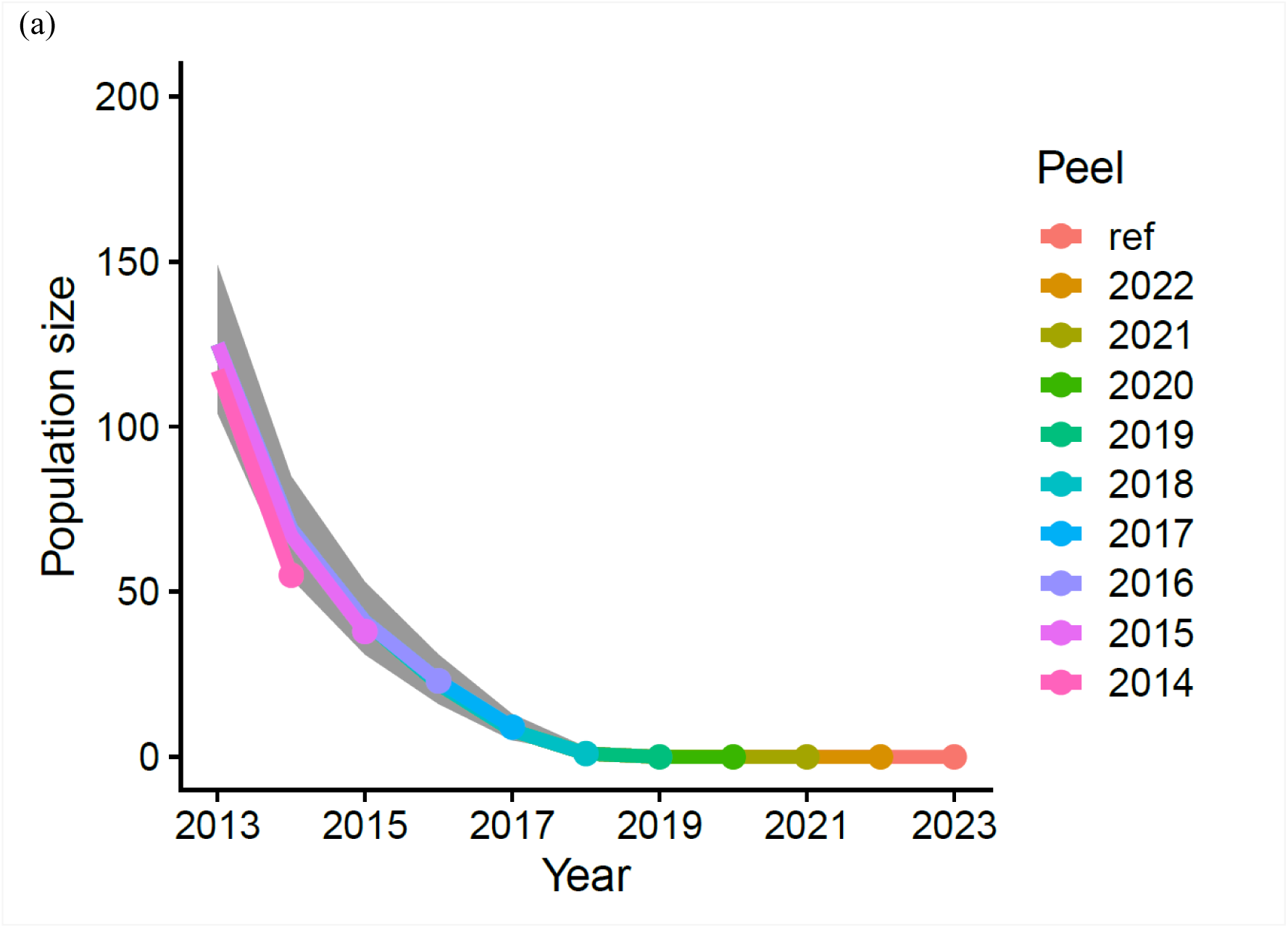
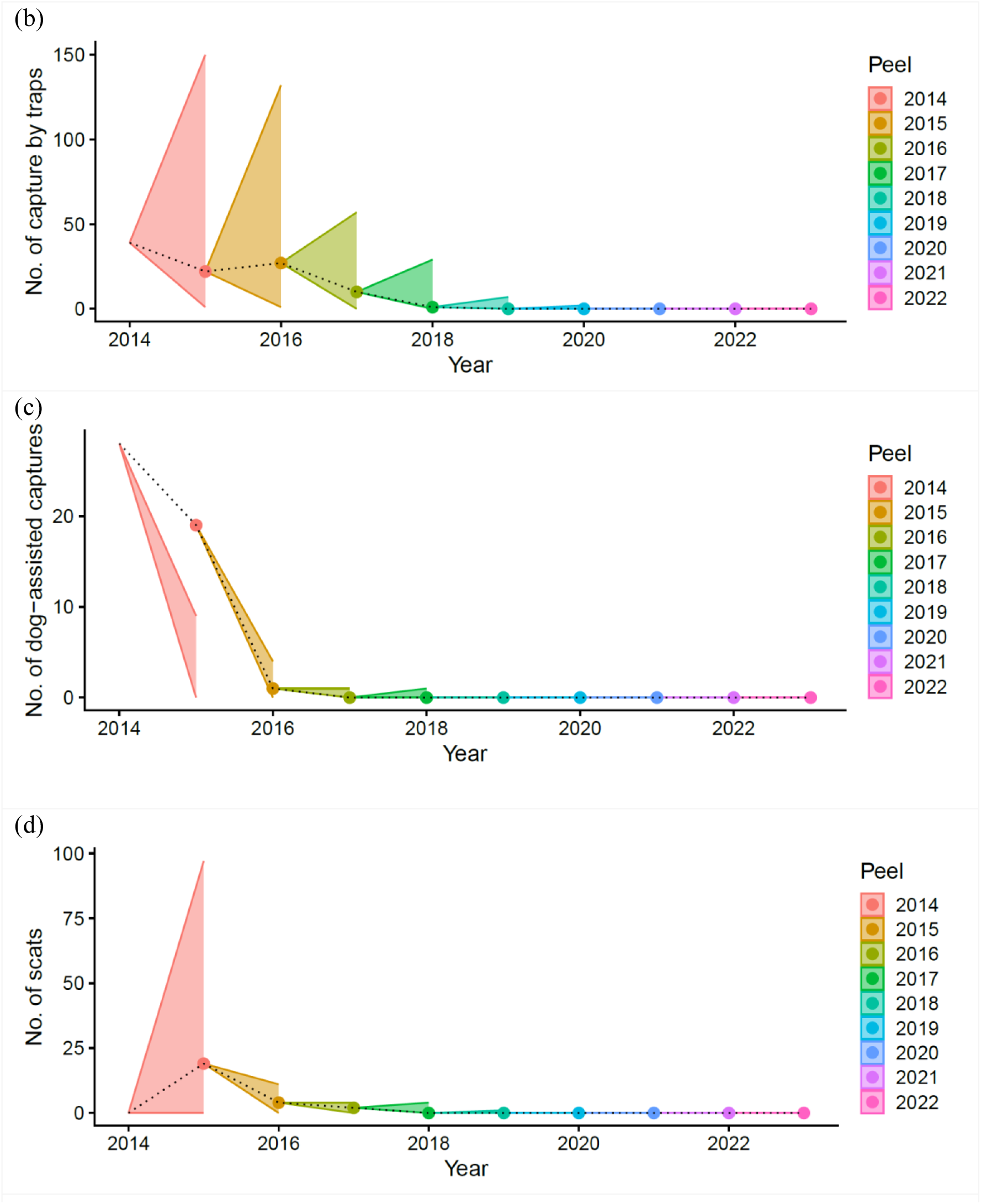
The results of retrospective analysis and hindcast cross varidation. (a) Retrospective analysis of population size estimates. The posterior medians estimated using datasets sequentially removed observations from the terminal year (peels) were shown on the reference posterior median using the full dataset (ref) and the 95% CI (shaded region). The hindcast prediction of three observation types: (b) number of captures by traps, (c) number of dog-assisted captures, (d) number of scats found by sniffer dogs. The shaded regions are 95% ranges of one-step-ahead predictions generated by every peels, and the dots and the dashed line are the true observations.

Eradication probabilities estimated by the hierarchical HBM were substantially higher than those obtained using the REA (Fig. 5). The HBM-based estimate increased to 0.953 in 2020 and reached 1.0 (numerically) in 2024 based on 10,000 posterior samples. In contrast, the REA estimated eradication probabilities of 0.744 in 2020 and 0.994 in 2024.

**Fig. 5.**
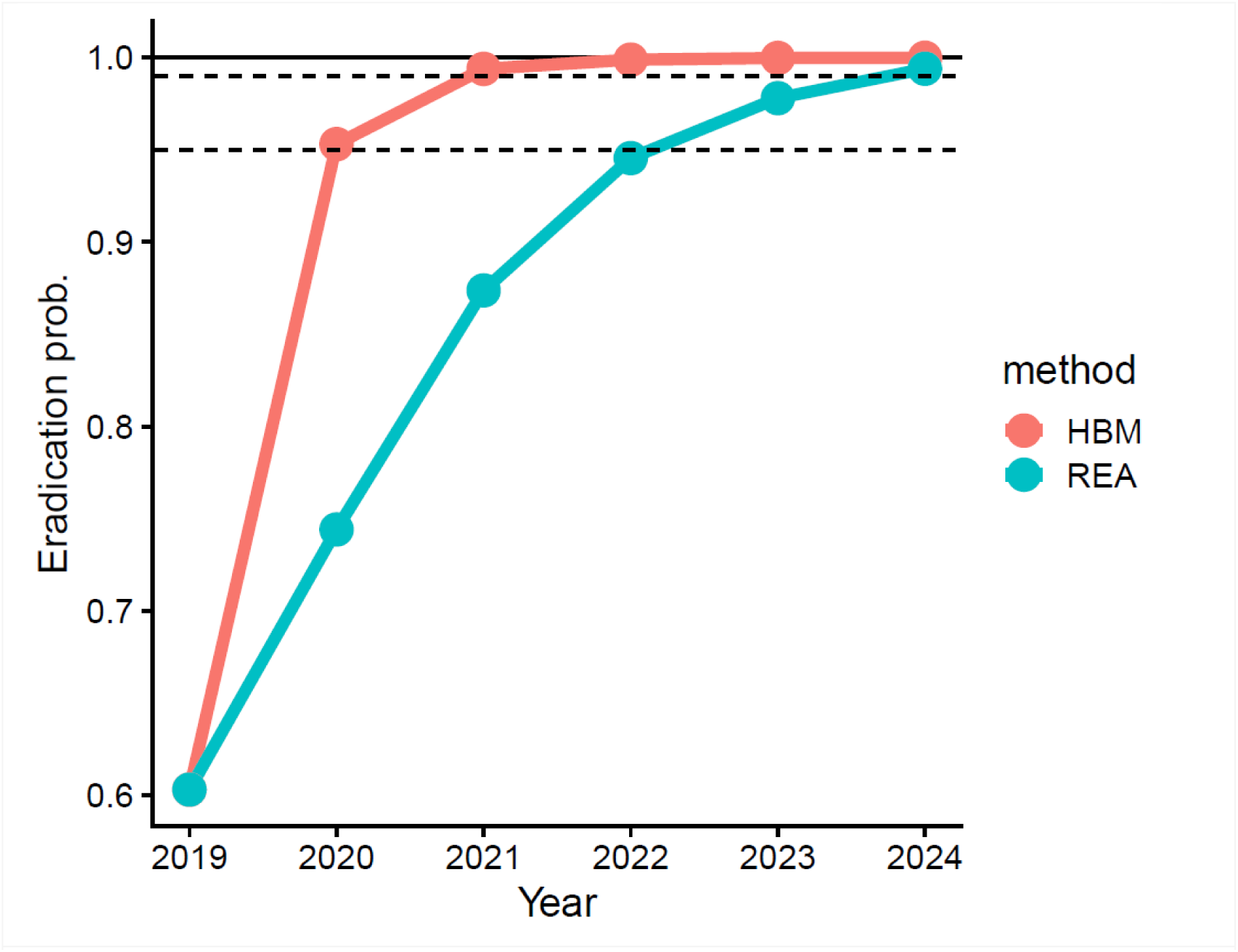
Eradication probability calculated by harvest-based model (HBM) and rapid eradication assessment (REA; Samaniego-Herrera et al., 2013, Appendix S2). Dashed lines indicate probabilities of 0.95 and 0.99.

## Discussion

Despite the recent trend toward eradication campaigns on large islands (Spatz et al., 2022), studies that document long-term trajectories of population size and removal rates leading to successful eradication remain limited. In this study, we reconstructed a 46-year trajectory of population size and capture rates for the small Indian mongoose using introduction records and long-term capture and monitoring data within a spatio-temporal hierarchical modeling framework. Population size estimates were robust to the addition of new data, and the predictive performance of the model improved as more data accumulated over time. Both the hierarchical HBM and REA indicated sufficiently high eradication probabilities by 2024, and no evidence of mongoose presence has been detected to date (June 2026). The reconstructed trajectories of population size and capture probability provide a comprehensive illustration of the dynamics underlying long-term, large-scale eradication programs. Furthermore, our approach provides interpretable indicators for evaluating management effectiveness and can be applied broadly to invasive species control programs.

Eradication feasibility generally depends strongly on detectability at low population densities and on meeting the condition that all individuals are exposed to removal efforts (“all individuals at risk” Bomford & O’Brien, 1995). In our case, mixture of passive traps and active dog-assisted capture methods might contribute to balance these criteria and maintain high catchablity until the removal of the last individual. A commonly noted principle in eradication programs is that the removal of the final fraction of a population can require disproportionately large effort compared to earlier stages. This highlights the importance of allocating sufficient resources over time to achieve complete eradication. In the Amami-Oshima program, the final 1% of individuals were removed after 2012. The caprure probability continued to rise since then (Fig. 3a), indicating that more annual capture effort has been expended than before. These results highlight the importance of sustained effort during the final stages of eradication.

The spatial allocation of removal effort is a key determinant of success in invasive species management (Baker, 2017; Nishimoto et al., 2020). In this case, control efforts initially concentrated near the introduction source and subsequently expanded outward across the island (Fig. 3b). Although the eradication program did not explicitly employ formal optimization methods, the observed allocation strategy qualitatively resembles theoretically optimal strategies for managing rapidly spreading populations (Baker, 2017). The delay between the peak of population size and the maximum invaded area (Fig. 2a; Fig. S3) is also consistent with predictions under optimal resource allocation. We note that the transition from a bounty-based system to a fully structured removal workforce in 2005 likely accelerated the decentralization in effort from the source area to the periphery, which would contributed to improve the efficiency of mongoose removal.

In long-term eradication programs, it is essential to update estimates of population status and management effectiveness as new data become available (Fukasawa, Hashimoto, et al., 2013). However, estimates that are overly sensitive to newly added data, or model predictions that are inconsistent with observations, can hinder decision-making (Carvalho et al., 2021). The low retrospective bias observed in our analysis (Fig. 4a; Fig. S5) indicates that the model provides stable estimates under data updates. Moreover, our population estimates were broadly consistent with those obtained from a previous non-spatial HBM (Fukasawa, Hashimoto, et al., 2013). The inclusion of both individual heterogeneity and spatio-temporal variation in catchability through the extended Weibull framework may have contributed to this stability. Model predictive performance improved over time (Fig. 4c, d; Fig. S6), highlighting the importance of accumulating sufficient data across all survey methods prior to declaring eradication.

Eradication probability is a useful metric for defining eradication success (Mazzamuto et al., 2020; Ramsey et al., 2009), but it is also sensitive to how the spatial structure of detection effort is represented (Samaniego-Herrera et al., 2013). Although the hierarchical HBM provides a powerful framework for reconstructing spatio-temporal population dynamics, its relatively coarse spatial resolution may lead to overestimation of eradication probability. This likely explains the higher estimates obtained from the HBM compared to the REA. We therefore recommend combining both approaches: HBM can provide estimates of key parameters such as population growth rate and prior eradication probability, which can then be used to inform more spatially explicit simulation-based assessments such as REA.

Eradication of invasive species on large islands remains challenging and typically requires adaptive management strategies to sustain high removal rates over extended periods. Hierarchical HBMs that fully exploit capture and monitoring data provide a powerful tool for reconstructing population trajectories, evaluating management performance, and predicting future outcomes. Although such models often require independent information on population size at some point in space or time (Fukasawa et al., 2020; Iijima, 2022; Kasada et al., 2023), recent advances in both marked (e.g., Fukasawa & Higashide, 2025) and unmarked (e.g., Nakashima et al., 2018) abundance estimation methods are making such data increasingly accessible. Wider application of these approaches, combined with robust monitoring and data archiving, will improve our ability to identify the key determinants of eradication success across systems.

## Acknowledgements

We express the deepest gratitude to the members of Amami Mongoose Busters for successful completion of the mongoose removal operations and development of massive dataset on capture history. This study was supported by the Environment Research and Technology Development Fund of the ERCA (JPMEERF20204006) funded by the Ministry of the Environment.

## Ethic Statement

The dataset used in this study except the spatial capture-recapture data belongs to the Japanese government and was collected in accordance with Japan’s laws, ordinances, and guidelines regarding animal welfare. Ethical approval for the spatial capture-recapture study using sniffer dogs was obtained from Okinawa University Animal Experimentation Committee (ID: 2020-03).

## Supporting Information

**Appendix S1** Additional figures.

**Appendix S2** Details of the simulation model applied to the rapid eradication assessment.

## Notes

### Competing Interest Statement

The authors have declared no competing interest.

