## Appendix S1 for "From introduction to eradication: reconstructing population size and removal history of an invasive species"

Appendix S1 Additional Figures

(a)

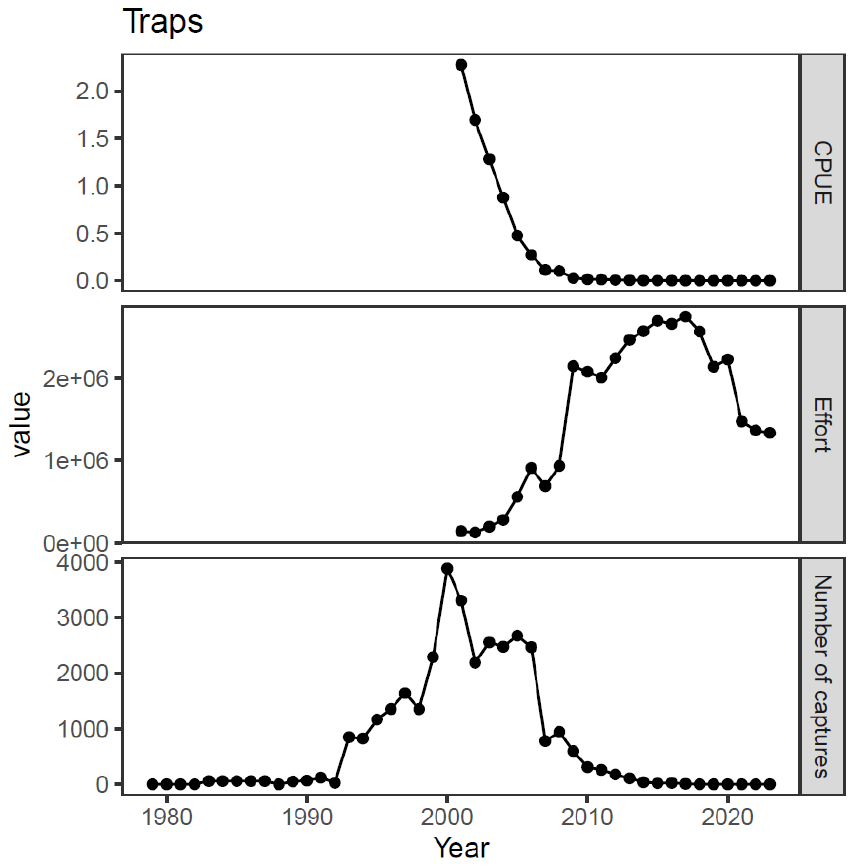

(b)

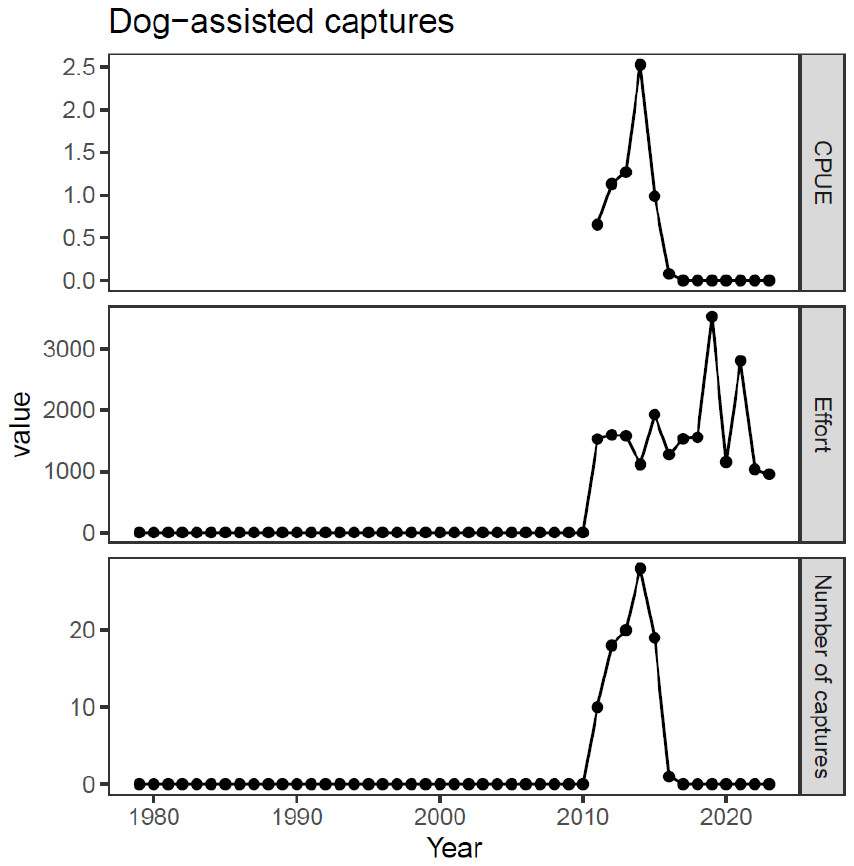

(c)

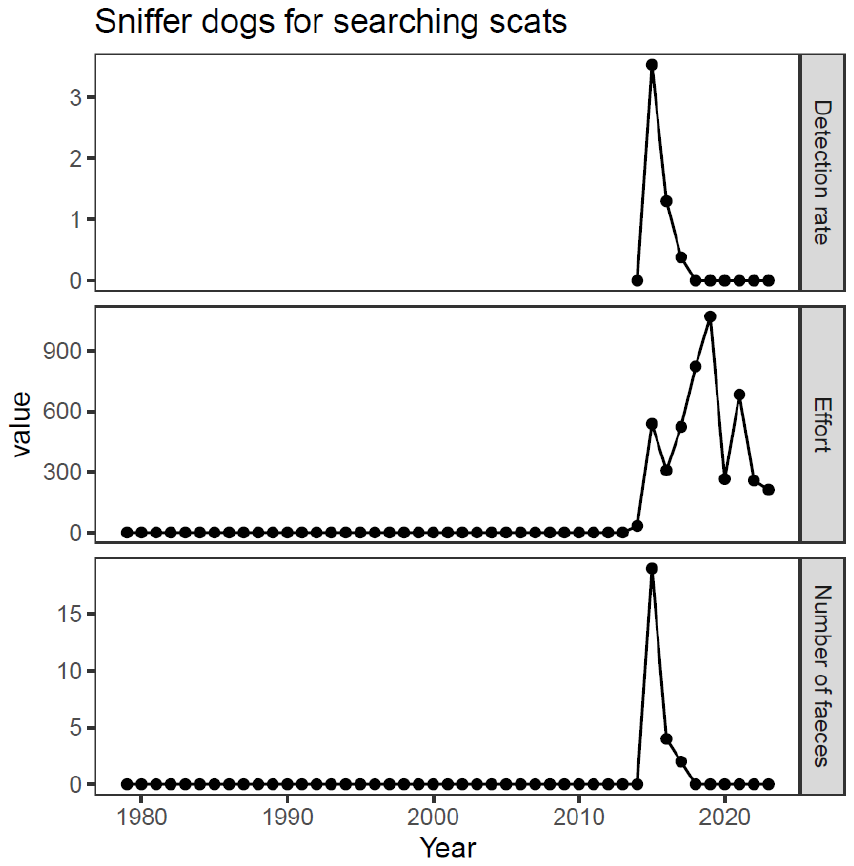

Fig. S1 The capture and monitoring dataset of mongoose removal. (a) Number of captures (detections), effort and catch (detections) per unit effort (CPUE) for traps, (b) dog-assisted captures, and sniffer dogs for searching scats. The units of effort were 100 corrected trap days for traps and search length (100 km) for dogs.

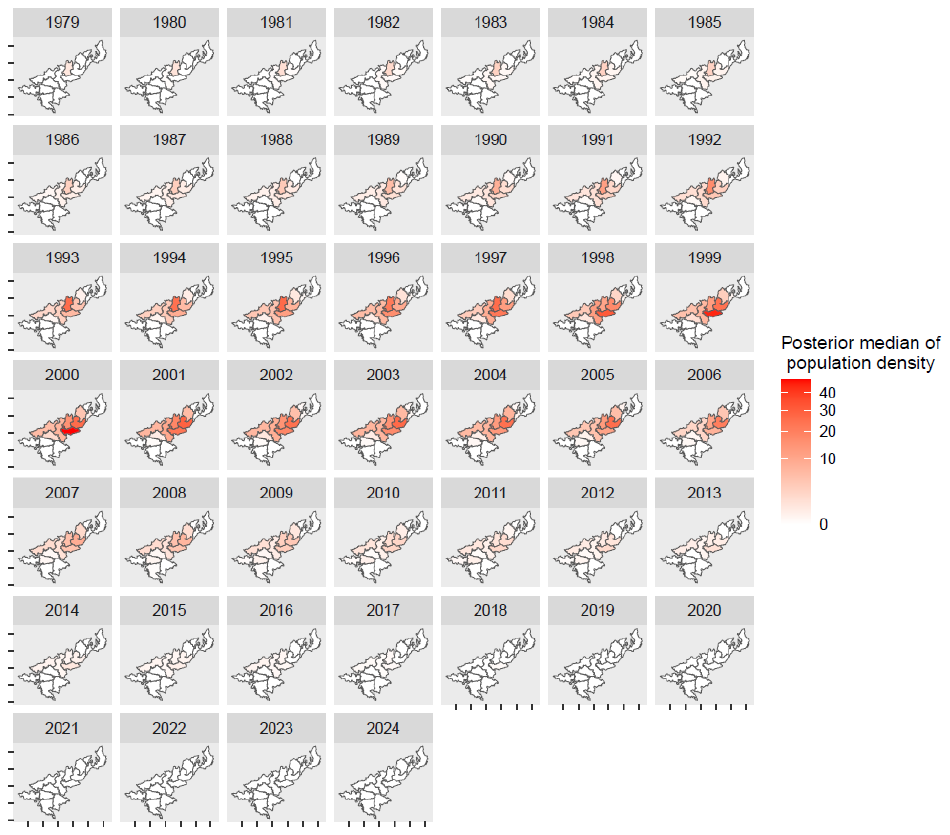

Fig. S2 Spatio-temporal patterns of the estimated population density.

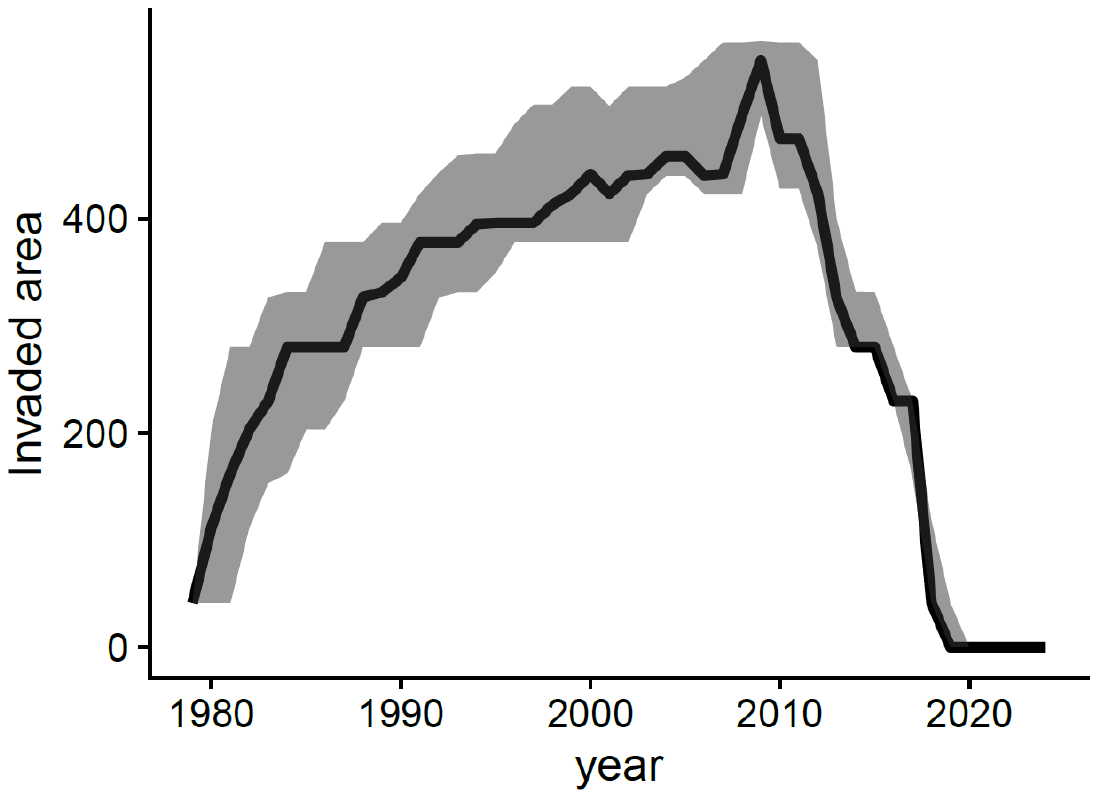

Fig. S3 Posterior median (solid line) and 95% CI of invaded area.

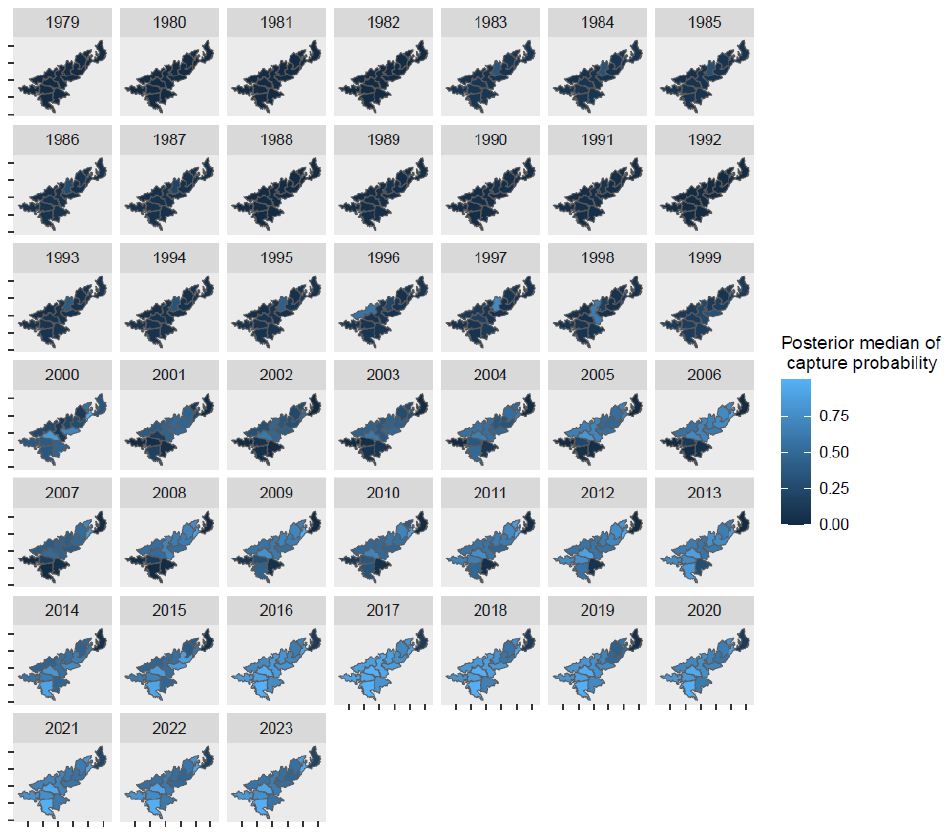

Fig. S4 Spatio-temporal patterns of capture probability.

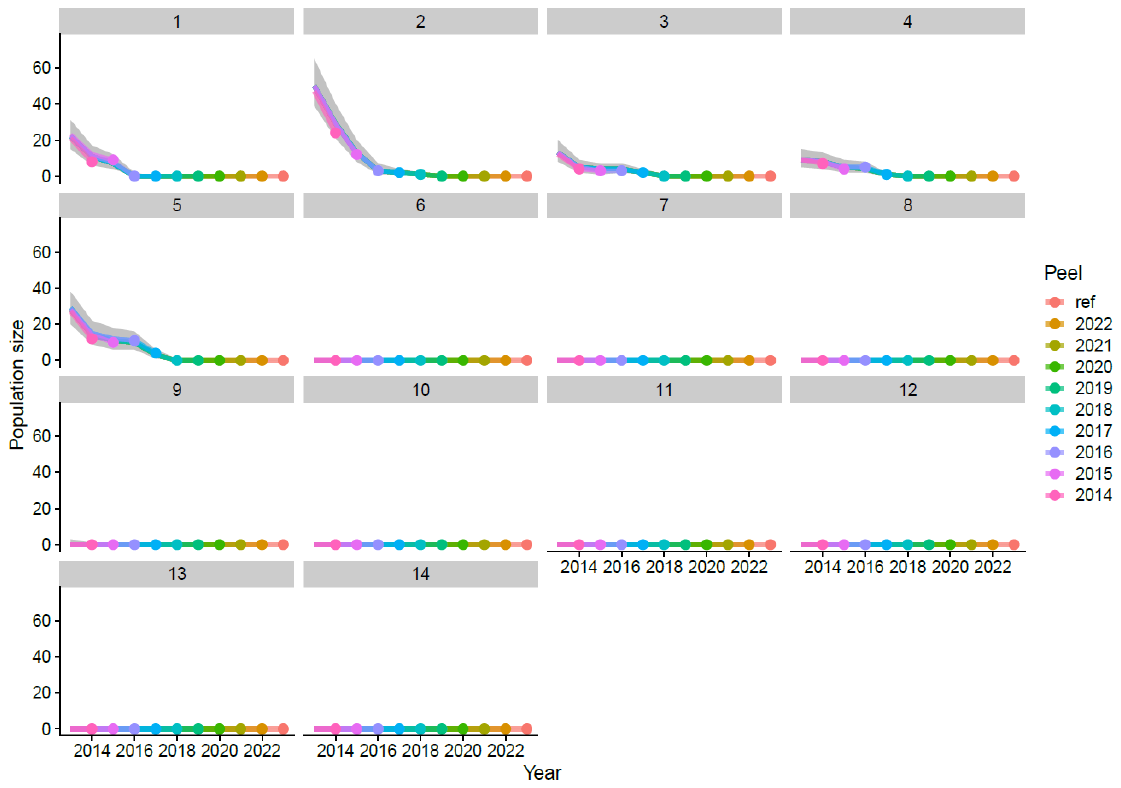

Fig. S5 The results of retrospective analysis of unit-level population size estimates. The posterior medians estimated using datasets sequentially removed observations from the terminal year (peels) were shown on the reference posterior median using the full dataset (ref) and the 95% CI (shaded region).

(a)

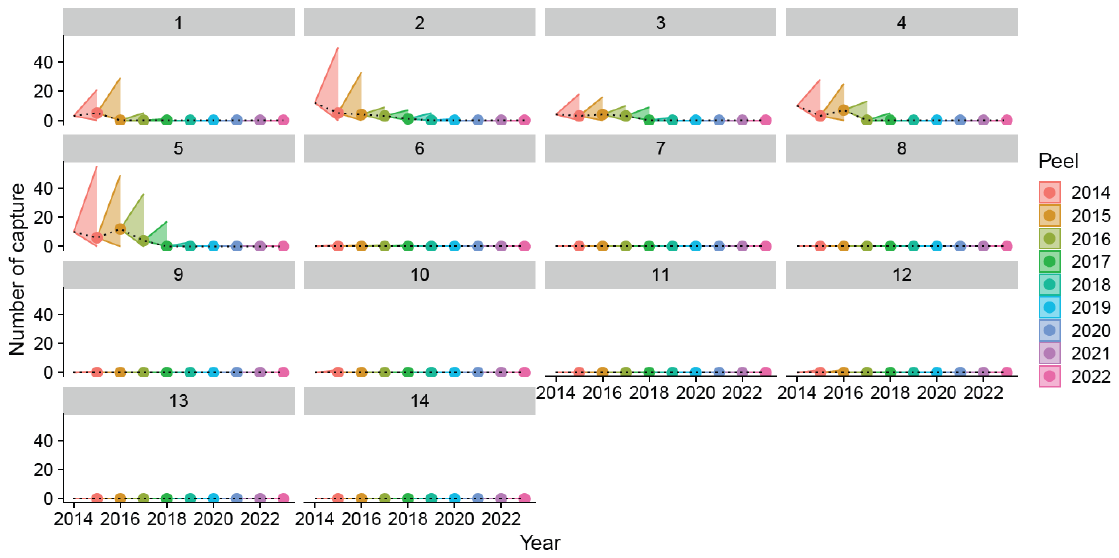

(b)

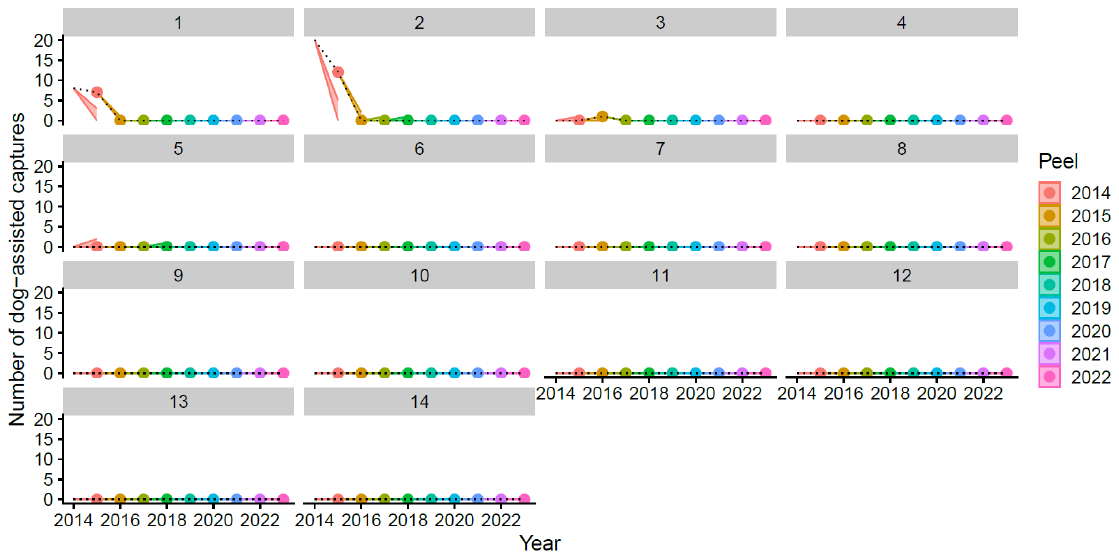

(c)

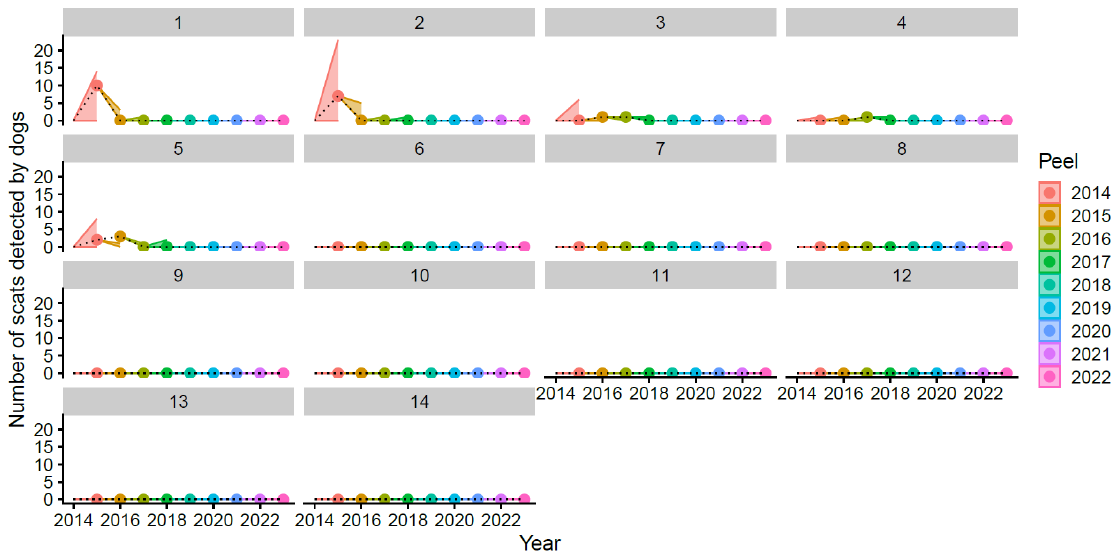

Fig. S6 The hindcast prediction of three observation types shown by units: (a) number of captures by traps, (b) number of dog-assisted captures, (c) number of faeces. The shaded regions are 95% ranges of one-step-ahead predictions generated by every peels, and the dots and the dashed line are the true observations.
