## Appendix S2 for "From introduction to eradication: reconstructing population size and removal history of an invasive species"

### Appendix S2 Details of the simulation model applied to the rapid eradication assessment

In this section, we describe the simulation methods used for the rapid eradication assessment (REA, Samaniego-Herrera et al., 2013) and the parameters applied in the simulations. The aim of the simulation is to estimate the detection probability in year *t* under the scenario that a pregnant survivor remains in 2019, the year following the last mongoose detection on the island. From a surveillance perspective, this represents a worst-case scenario, as detecting a single individual is more difficult than detecting multiple individuals.

The simulation is individual-based and spatially explicit. In each iteration, the following steps were performed: (1) random determination of the activity center of the surviving individual, (2) reproduction and dispersal of offspring, (3) detection of individuals by survey methods (i.e., traps and dogs), and (4) repetition of steps (2) and (3) until 2024. Most parameters used in the simulation were derived from the harvest-based model (HBM) or from existing literature, while parameters related to home range size and detectability were estimated using a DNA-based capture–recapture survey.

Reproduction and dispersal of offspring were modeled stochastically. The number of offspring produced by an individual was drawn from a Poisson distribution with mean equal to intrinsic growth rate. The spatial location of each offspring was represented by its activity center,, **x***_offspring_*, which was sampled from a two-dimensional Gaussian distribution centered on the parent’s location, **x***_parent_*:
**x***_offspring_* ~ N_2_(**x***_offspring_*| **x***_parent_*, *σ*^2^**I**)
The initial location of the surviving individual was randomly assigned within the island. To avoid unrealistic placement in areas where mongoose presence was highly unlikely, we used posterior probabilities (estimated by the HBM) that population size was non-zero in 2019 as sampling weights across management units.

To simulate the detection process, the spatial configuration of traps and the trajectories of dog surveys were reconstructed within the simulation space. Detection effort was represented as point locations in space, each associated with an annual effort. Detection efforts of traps and dog-assisted capture were standardized by scaling their observed detection rates per unit effort so that they were comparable to the search distance of scat-detection dogs. Dog survey trajectories were discretized by extracting edges from GPS tracking data.

The detection probability of individual *i* by a unit of detection effort *j*, denoted *p_ij_*, depends on the Euclidean distance between the individual’s activity center and the location of the detection effort, *Z_ij_* ,and the amount of effort, *E_j_* :
$p_{ij}=1-{{(1-g}_{0}exp(-\frac{{Z_{ij}}^{2}}{2\tau^{2}}))}^{E_{j}}$

where g_0_ is the baseline detection probability at the activity center, and τ determines the spatial scale of detection (i.e., home range size). Detection of individual *i* across all effort points was determined by a Bernoulli trial with probability:
 1 - Π*_j_*(1 - *p_ij_*)

From each simulation iteration, we recorded whether at least one mongoose was detected in year *t*. A total of 10,000 iterations were conducted, and detection probability was estimated as P(detection)*ₜ*.

Parameters used in the simulations are summarized in Table S1. To account for parameter uncertainty, values were sampled from their respective distributions in each iteration. The posterior distribution of the HBM was used for the population growth rate, approximated by a normal distribution on the log scale. The dispersal parameter (σ) of the Gaussian kernel was derived from the reported invasion front speed of mongoose (600-1000 m per year; Abe et al., 1991), , following Skellam (1951)’s diffusion framework. In this framework, the diffusion coefficient *D* is related to invasion speed *v* and population growth rate *r* as:
*D* = (2*v*)^2^*r*.
The variance of the Gaussian dispersal kernel is given by *σ*^2^ = 2*D*.

Table S1 Parameters used in the simulation, their disribution and source. All the distributions applied were two-parameter distributions, and the column *p*_1_ and *p*_2_ are the parameter values in the distributions.

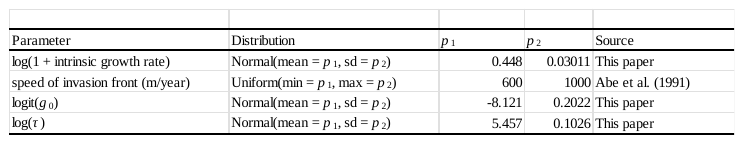

We conducted a faecal DNA mark-recapture study to estimate *g*_0_ and *τ*. To this end, we carried out a scat-detection survey using the same protocol as that employed in the scat-detection dog monitoring program of the mongoose control campaign, and collected faecal samples for individual identification based on DNA analysis. The field survey was conducted 24 days between December 2020 and May 2021 in forested areas and along roads within Nago City on Okinawa Island, where mongoose populations are well established. A total survey distance of 180.51 km resulted in the collection of 726 faecal samples.

We extracted DNA from the faecal samples using QIAamp Fast DNA Stool Mini Kit (QIAGEN, Valencia, California, USA). To confirm the presence of mongoose DNA in the samples, PCR and electrophoresis were performed using mongoose-specific primers targeting 340 bp of the mitochondrial cytochrome b gene (Imazato et al., 2012). PCR with the species-specific primers was conducted up to two times, and samples that failed to amplify were excluded from further analyses. For individual identification, we used 11 simple sequence repeat (SSR) markers (Uau2, Uau3, Uau4, Uau8, Uau9, Uau10, Uau11, Uau13, Uau14, Uau15, and Uau17) and one sex-identification marker developed by Sato et al. (2021). To improve experimental efficiency, we modified the method of Sato et al. (2021) by directly labeling the 5′ ends of the forward primers with four different fluorescent dyes and using them in multiplex PCR. Following Lampa et al. (2013), heterozygous and homozygous genotypes were accepted only when confirmed by two and three independent PCR replicates, respectively. Consensus genotypes were then determined, and samples with missing data at more than four of the 12 loci were excluded from further analyses. Individual identification was conducted in GIMLET (Valière, 2002) using the Regroup option, and the probability of identity (PID) and sibling probability of identity (PIDsib) were calculated. In total, 139 samples from 95 individuals were retained. The PID and the PIDsib were 1.40 × 10^−6^ and 1.49 × 10^−3^, respectively.

The parameters *g*_0_ and *τ* were estimated using a spatially explicit capture-recapture model (SECR, Borchers & Efford, 2008). We discretized GPS tracks into vertices and allocated the distances traveled along each track segment to the corresponding vertices. In this way, each vertex represents a survey location associated with a measure of effort, allowing it to be treated as both the position and magnitude of survey effort within the SECR framework. The numbers of genotyped samples for each individual were also assigned to the corresponding vertices.

A buffer of 1,000 m was added around the bounding box of surveyed locations for numerical integration of activity centers; this buffer size is sufficiently large relative to the home-range size of mongoose. Parameters were estimated using an SECR model with a Poisson detection function, in which detection probability was adjusted by distance traveled as a measure of spatially varying effort, and a Gaussian detection kernel was assumed for home-range structure. Maximum likelihood estimation was conducted using the function secr() in the R package secr 5.1.0 (Efford, 2022). The estimate of *g*_0_ obtained from the SECR model corresponds to the detectability of successfully genotyped samples. To obtain the detectability of all mongoose faecal samples, we adjusted this estimate by accounting for the genotyping success rate.
